# Molecular architecture and assembly of the human RNA Degradosome

**DOI:** 10.64898/2026.09.03.749114

**Authors:** Justin A. Gee, Jason G. Williams, Monica C. Pillon

## Abstract

The PNPase exoribonuclease interacts with the SUV3 helicase to form the human RNA Degradosome responsible for processive mitochondrial RNA turnover and quality control. Although structural data reveals the trimeric PNPase architecture along with the monomeric and dimeric states of SUV3, the molecular basis of their assembly for processive RNA degradation remains poorly understood. Here we present the molecular characterization of the human RNA Degradosome and define critical molecular features that regulate complex formation. We identify the protein regions required for RNA Degradosome formation and establish the minimum components necessary for the SUV3-PNPase interaction. Chemical crosslinking mass spectrometry analysis reveals distance restraints that position the SUV3 N-terminal domain alongside of the PNPase S1 domain, defining an important interface for complex formation. These findings provide mechanistic insight into how PNPase and SUV3 assemble to form the human RNA Degradosome.

## Introduction

The RNA Degradosome maintains cellular homeostasis through mitochondrial RNA degradation (Wang et al. 2010; Borowski et al. 2013; Dhir et al. 2018; Liu et al. 2018). The human RNA Degradosome is a multi-enzyme complex comprised of the PNPase 3’→5’ exoribonuclease and the SUV3 ATP-dependent helicase (Borowski et al. 2013). Together, they mediate the processive turnover of most mitochondrial RNAs to support RNA surveillance and post-transcriptional regulation of essential oxidative phosphorylation genes (Pietras et al. 2018; McShane et al. 2024). In the absence of SUV3, mitochondrial RNAs are largely resistant to PNPase-mediated degradation, leading to the accumulation of deleterious RNA intermediates (Wang et al. 2009; Szczesny et al. 2010). The importance of mitochondrial RNA degradation is underscored by clinical mutations in the PNPase gene (PNPT1), which can cause mitochondrial dysfunction and severe neurodevelopmental disorders such as Leigh Syndrome (Vedrenne et al. 2012; von Ameln et al. 2012; Alodaib et al. 2016; Matilainen et al. 2017; Golzarroshan et al. 2018; Sato et al. 2018).

The multidomain PNPase exoribonuclease relies on intrinsic flexibility for RNA degradation. PNPase adopts a stable homotrimeric structure in which the RNase PH1, α-helical, and RNase PH2 domains assemble into a barrel with a narrow central channel (Lin et al. 2012; Jain et al. 2022). This channel runs the length of the barrel and selectively accommodates unstructured RNA, with flexible loops directing the threaded RNA towards the opposite end of the barrel, where the exoribonuclease active sites mediate substrate degradation (Unseld et al. 2025). The PNPase C-terminus comprises the KH domain, S1 domain, and a flexible tail that lines the entrance of the channel, where it contributes to substrate binding (Lin et al. 2012; Unseld et al. 2025). Recent structural work reveals the extensive conformational flexibility of the S1 domains, adopting distinct open and closed states at the channel entrance important for regulating substrate capture and degradation (Li et al. 2025).

The asymmetric SUV3 homodimer functions as a processive RNA helicase. SUV3 is a multidomain helicase comprised of an N-terminal domain (NTD), tandem RecA1 and RecA2 domains, a C-terminal domain (CTD), and a flexible tail (Jedrzejczak et al. 2011). Although the functional roles of the NTD and CTD remain underexplored, the RecA1 and RecA2 domains form the SF2 helicase core with a nucleotide binding site located at its interface to couple ATP hydrolysis with RNA unwinding and translocation (Jedrzejczak et al. 2011). Human SUV3 adopts a concentration-dependent monomer-dimer equilibrium, where the higher-order oligomer exhibits enhanced RNA processivity (Jain et al. 2022). The asymmetric SUV3 homodimer positions its two protomers 110° relative to one another, with only one protomer bound to both nucleotide and RNA substrate (Jain et al. 2022; Patra et al. 2026). The dimeric protomers are functionally distinct enabling one subunit to catalyze RNA unwinding and translocation, while the second acts allosterically to stimulate rapid ATP hydrolysis to prevent RNA dissociation (Patra et al. 2026).

Although SUV3 and PNPase are known to assemble into a functional RNA Degradosome, the molecular basis for their association remains largely unclear. The SUV3 dimer directly associates with the PNPase trimer to form a large hetero-pentameric complex that efficiently degrades double-stranded RNA in an ATP-dependent manner (Wang et al. 2009; Jain et al. 2022). Deletion of the SUV3 flexible tail, which is required for homodimerization, disrupts PNPase association *in vitro* suggesting the oligomer may be required for complex formation (Jain et al. 2022). However, the SUV3 monomer retains intermediate stimulation of PNPase RNA degradation activity, indicating that it may still associate with PNPase, albeit less efficiently (Jain et al. 2022). An early biochemical study also reports that a short deletion within the SUV3 CTD disrupts PNPase binding (Wang et al. 2009). However, subsequent SUV3 structures suggests that this deletion may alter the local structure of the CTD, raising the possibility that its effect on complex formation could be either direct or indirect (Jedrzejczak et al. 2011; Patra et al. 2026). Recent characterization of disease-associated PNPase variants identified the D713Y mutation as deficient in SUV3 binding, suggesting that the conformationally dynamic C-terminal region of PNPase may contribute to RNA Degradosome assembly (Li et al. 2025). This is supported by small angle X-ray scattering data of the RNA Degradosome in which the SUV3 dimer is suggested to cap the channel entrance of the PNPase exoribonuclease (Jain et al. 2022). Although these studies have provided important insight into the overall architecture of the human RNA Degradosome, the molecular organization and interaction interfaces governing complex assembly remains poorly understood.

To define the molecular basis of RNA Degradosome assembly, we generated a series of SUV3 and PNPase variants to define the requirements for complex formation. Using affinity pulldown assays, chemical crosslinking mass spectrometry analysis, and molecular modeling, we identify an interaction between the SUV3 NTD and the dynamic PNPase S1 domain that positions the helicase at the entrance of the ribonuclease channel. Collectively, these findings provide new insight about the SUV3-PNPase interface and support a model for how SUV3 engages PNPase for processive degradation of mitochondrial RNA.

## Results

### SUV3 NTD is required for PNPase association

To evaluate the molecular determinants of the SUV3-PNPase interaction, we performed an affinity pulldown assay of the human RNA Degradosome from mammalian cell lysate. We transiently transfected HEK293F cells with plasmid DNA encoding either C-terminally tagged human SUV3-Strep or human PNPase-Flag. Western blot analysis confirmed selective capture of SUV3-Strep by StrepTactin due to its strong enrichment in the resin bound fraction, whereas PNPase-Flag was not detectable (Figure 1). Next, SUV3-Strep was transiently transfected along with PNPase-Flag. Affinity purification of SUV3-Strep in physiological salt conditions resulted in the co-purification of PNPase-Flag, confirming previous reports that the human SUV3 helicase and PNPase exoribonuclease undergo a detectable interaction (Figure 1) (Wang et al. 2009; Jain et al. 2022).

**Figure 1.**
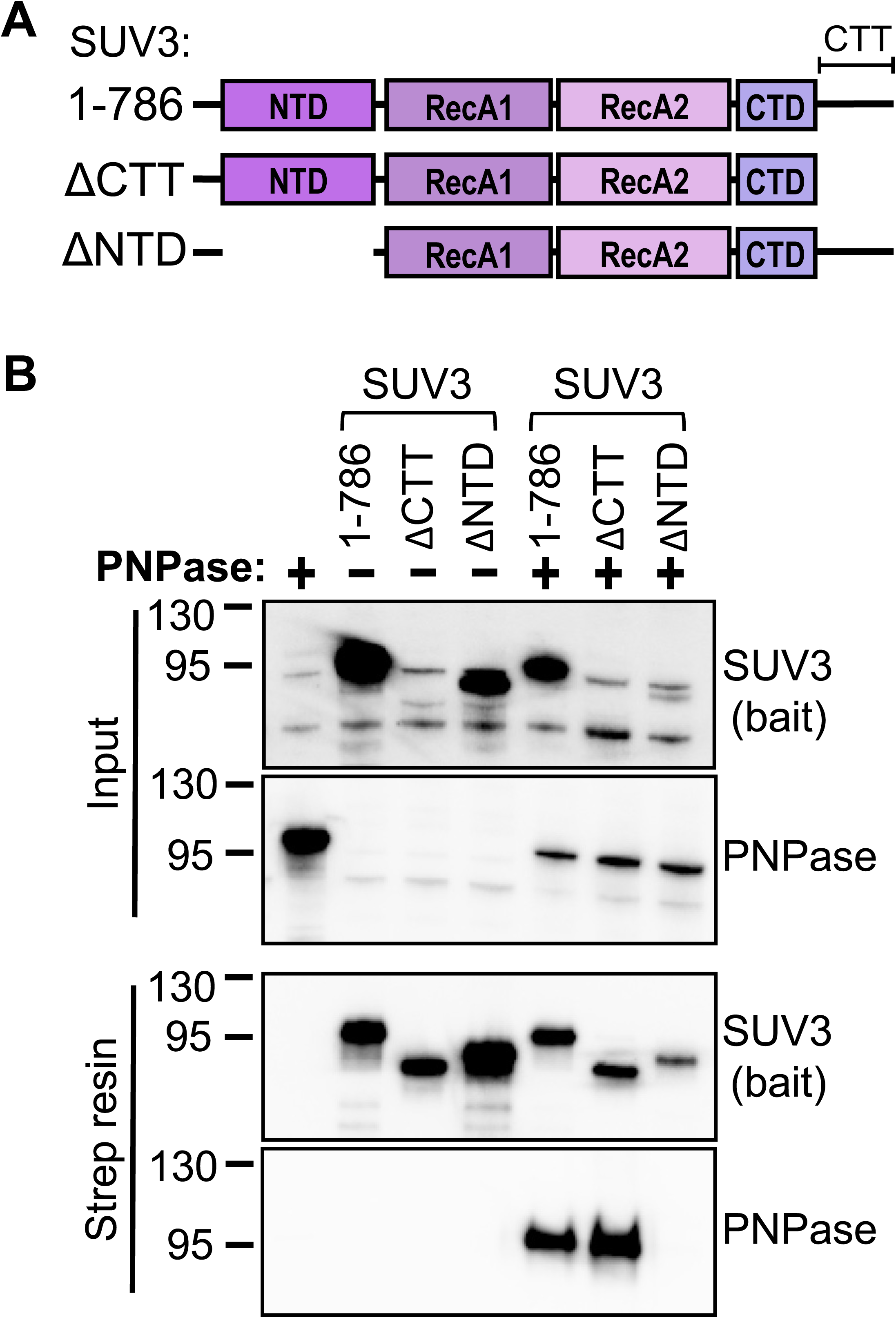
The SUV3 NTD is indispensable for PNPase association. *A*, Domain architecture of human SUV3 truncations. The N-terminal domain (NTD) is dark purple, the RecA1 domain is light purple, the RecA2 domain is pink, the C-terminal domain (CTD) is violet, and the C-terminal tail (CTT) is marked by a bracket. *B*, Representative co-immunoprecipitation and Western blot analysis of transiently transfected SUV3-TwinStrep truncations along with full-length PNPase-3xFlag. SUV3 variants were immobilized on the anti-Strep affinity support, serving as the bait. The SUV3 NTD is required to retain PNPase-3xFlag on the anti-Strep affinity support. Input defines the clarified whole cell lysate.

To define the molecular features of SUV3 required for RNA Degradosome assembly, we monitored complex formation using a series of SUV3 truncation variants. We generated two SUV3 truncations by either removing the NTD or the flexible C-terminal tail (CTT), both of which have been implicated in SUV3 dimerization (Jain et al. 2022; Patra et al. 2026). Despite the removal of the CTT and the resulting impairment of SUV3 dimerization (Jain et al. 2022), the SUV3^ΔCTT^-Strep variant retained RNA Degradosome assembly, indicating that both the CTT and SUV3 dimerization are dispensable (Figure 1). Conversely, deletion of the NTD (SUV3^ΔNTD^) prevented the co-purification of PNPase-Flag, indicating a loss in PNPase association (Figure 1). Together, this analysis demonstrates that RNA Degradosome assembly is dependent on the SUV3 NTD, but not the CTT or helicase dimerization.

### PNPase S1 domain is essential for RNA Degradosome assembly

Next, we performed a reciprocal experiment to characterize the SUV3-PNPase interaction using a series of engineered PNPase truncations. We systematically removed elements forming the entrance of the PNPase RNA channel, including the KH domain, S1 domain, and the flexible tail (Figure 2A). These PNPase variants were transiently expressed and co-purified with SUV3-Strep using StrepTactin. Deletion of the flexible tail (PNPase^1-753^) had no appreciable effect on SUV3 binding properties indicating that the PNPase tail is not required for RNA Degradosome assembly (Figure 2B). Conversely, removal of the PNPase S1 domain (PNPase^1-673^) abolished SUV3 association (Figure 2B). Likewise, deletion of the entire PNPase C-terminus (PNPase^1-598^), beginning at the KH domain, abrogated the association with SUV3. Together, these data reveal that the PNPase S1 domain is indispensable for RNA Degradosome assembly and that the entrance of the RNA channel harbors an important SUV3 binding interface.

**Figure 2.**
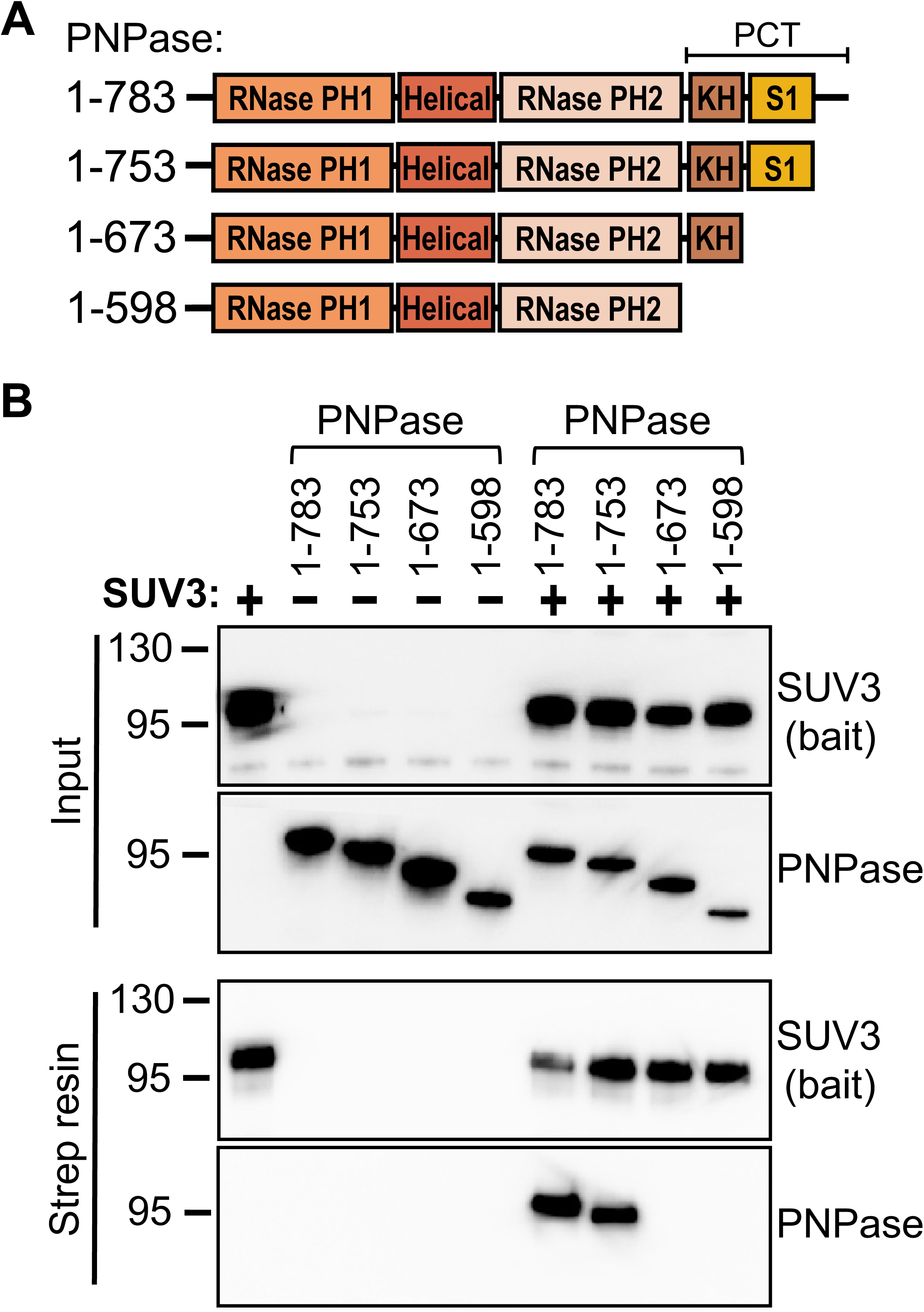
The PNPase S1 domain is essential for RNA Degradosome assembly. *A*, Domain architecture of human PNPase truncations. The RNase PH1 domain is light orange, the helical domain is dark orange, the RNase PH2 domain is beige, the KH domain is brown, and the S1 domain is gold. The PNPase C-terminus (PCT) comprises the KH domain, S1 domain, and labile tail and is defined by a bracket. *B*, Representative co-immunoprecipitation and Western blot analysis of transiently transfected full-length SUV3-TwinStrep along with PNPase-3xFlag truncations. SUV3 was immobilized on the anti-Strep affinity support, serving as the bait. The PNPase S1 domain is necessary for its association with SUV3. Input defines the clarified whole cell lysate.

### A single PNPase subunit is sufficient for SUV3 binding

After establishing the importance of the S1 domain, we determined whether the PNPase C-terminus is sufficient for RNA Degradosome assembly. We substituted the PNPase barrel for a ring-shaped molecular scaffold of comparable diameter. The Flag-tagged molecular scaffold exploits the architectural similarity between the PNPase barrel and the eukaryotic DNA clamp PCNA to display three copies of the PNPase C-terminus (^3x^PCT), comprising the KH domain, S1 domain, and the flexible tail, one a single face of the ring (Figure 3A). The ^3x^PCT conjugated scaffold co-purified with SUV3-Strep whereas a control scaffold (^3x^CTL) lacking the PNPase moieties failed to association with SUV3 (Figure 3B). This experiment confirms that the PNPase C-terminus is sufficient for facilitating an SUV3 interaction.

**Figure 3.**
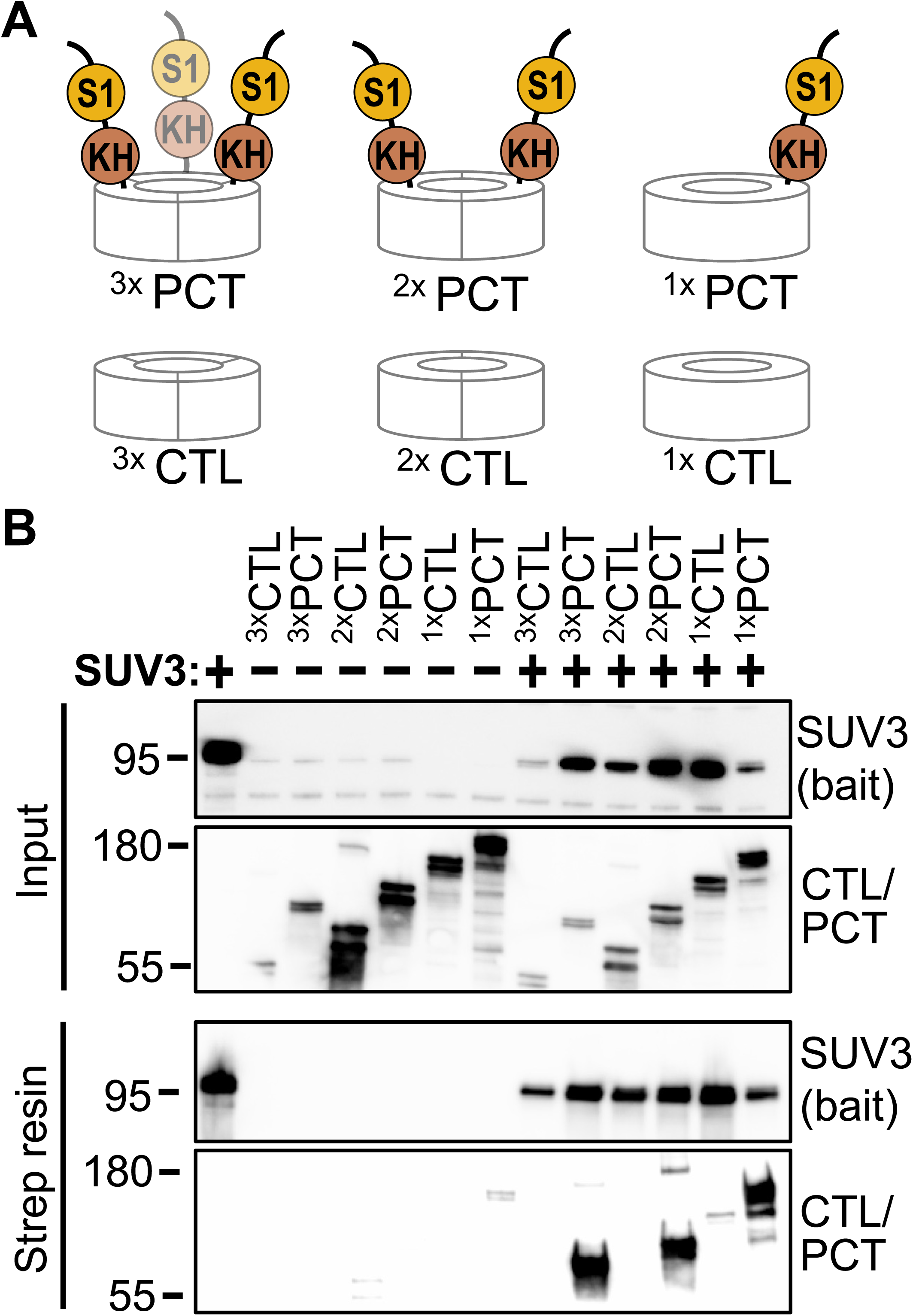
The PNPase C-terminus is sufficient for SUV3 binding. *A*, Cartoon schematic of an adaptable ring scaffold presenting three equivalent termini (3x), two equivalent termini (2x), or a single terminus (1x). Control scaffolds (CTL) lack a C-terminal PNPase moiety, whereas the modified scaffolds display the PNPase C-terminus (PCT) comprising the KH domain, S1 domain, and labile tail. *B*, Representative co-immunoprecipitation and Western blot analysis of transiently transfected full-length SUV3-TwinStrep along with scaffold ring variants. SUV3 was immobilized on the anti-Strep affinity support, serving as the bait. A single PCT moiety is sufficient to bind SUV3. Input defines the clarified whole cell lysate.

We next sought to determine whether all three subunits of the PNPase C-terminus are required for SUV3 association. To control for the number of PNPase moieties presented on the ring, we varied the number of termini by altering the rotational symmetry of the molecular scaffold. PCNA is a homotrimer with three evenly spaced termini, whereas the structurally homologous bacterial β clamp is a homodimer with two juxtaposed termini, and an engineered β clamp–β clamp’ chimeric ring has a single terminus (Figure 3A). Control scaffolds were outfitted to display the PNPase C-terminal region at their corresponding stoichiometry. In the absence of the PNPase C-terminal moieties, all three control scaffolds lacked binding affinity for SUV3-Strep, indicating the scaffolds are inert in the conditions tested (Figure 3B). On the other hand, scaffolds presenting three, two, and even one copy of the PNPase C-terminus exhibited robust SUV3 binding properties. This analysis indicates that a single copy of the PNPase C-terminus is sufficient for SUV3 association.

### Chemical crosslinking mass spectrometry analysis of SUV3-PNPase

To further investigate the human RNA Degradosome assembly, we reconstituted the SUV3-PNPase complex at a 2:3 molar ratio, consistent with the reported stoichiometry of the machinery (Wang et al. 2009). We performed an *in vitro* RNA degradation assay to confirm the association between purified SUV3 and PNPase enzymes. Complex assembly was assessed by monitoring degradation of a 3′-overhang RNA duplex substrate previously used to characterize RNA Degradosome activity (Wang et al. 2009). As expected, PNPase alone did not exhibit detectable RNA degradation activity in the conditions tested, recapitulating earlier observations that the exoribonuclease is not capable of driving degradation of duplexed RNA (Supplementary Figure S1) (Wang et al. 2009). Purified SUV3 also lacked ribonuclease activity, confirming the sample was sufficiently pure of trace nucleases (Supplementary Figure S1). On the other hand, the mixture of PNPase with the SUV3 helicase licensed the exoribonuclease for RNA degradation (Supplementary Figure S1). The observed exoribonuclease activity was attributed to PNPase as SUV3 failed to stimulate the nuclease-deficient PNPase D544N variant (Supplementary Figure S1). Together, this data confirms successful *in vitro* reconstitution of the human RNA Degradosome complex.

To define the structural organization of the RNA Degradosome assembly, we applied chemical crosslinking mass spectrometry (XL-MS). We used the amine-reactive chemical crosslinker bis(sulfosuccinimidyl)suberate to determine RNA-independent spatial constraints between the SUV3 helicase and PNPase exoribonuclease. The crosslinker was titrated into the SUV3-PNPase mixture to optimize crosslinking efficiency of surface accessible primary amines, such as lysine residues or the protein’s N-terminus. Chemical crosslinks were identified under reaction conditions that enriched discrete crosslinked species while minimizing nonspecific smearing observed by denaturing gel electrophoresis (Supplementary Figure S2). Mass spectrometry analysis revealed approximately 265 unique residue-residue crosslink pairs, including high quality spectra supporting SUV3-SUV3, PNPase-PNPase, and SUV3-PNPase interactions (Supplementary Figure S3).

We detected chemical crosslinks that support the structural architecture of the previously reported SUV3 homodimer and PNPase homotrimer. We observe extensive crosslinking across the SUV3 CTT (Figure 4A). This broad spatial crosslinking pattern is consistent with a conformationally flexible CTT, which is supported by the lack of Coulomb potential density for the CTT in the recent SUV3 cryoEM reconstruction (Patra et al. 2026). Most SUV3 crosslinks involved the labile CTT, preventing modeling onto available experimental SUV3 structures. The remaining crosslinks were mapped onto the reported asymmetric SUV3 dimer bound to both single-stranded RNA and nucleotide (Patra et al. 2026). The 18 unique intra-molecular SUV3 crosslinks were compatible with the canonical SF2 helicase protein fold whereas 3 unique inter-molecular crosslinks are consistent with the asymmetric dimeric arrangement (Figure 4B) (Jedrzejczak et al. 2011; Patra et al. 2026). We also observed broad crosslinking coverage across the PNPase enzyme, consistent with its homotrimeric fold (Figure 4C). Unique intra-molecular (total 12) and inter-molecular (total 8) crosslinks support the canonical fold and arrangement of the PNPase barrel and pore forming KH domains (Figure 4D) (Lin et al. 2012; Unseld et al. 2025). The reported open and closed PNPase states adopted through asymmetric S1 domain conformational changes prevents confident assignment of S1 domain crosslinks to a single static structure (Figure 4E) (Li et al. 2025). Therefore, additional crosslinks within the S1 domains support the domain fold but could not be unambiguously assigned as inter-molecular.

**Figure 4.**
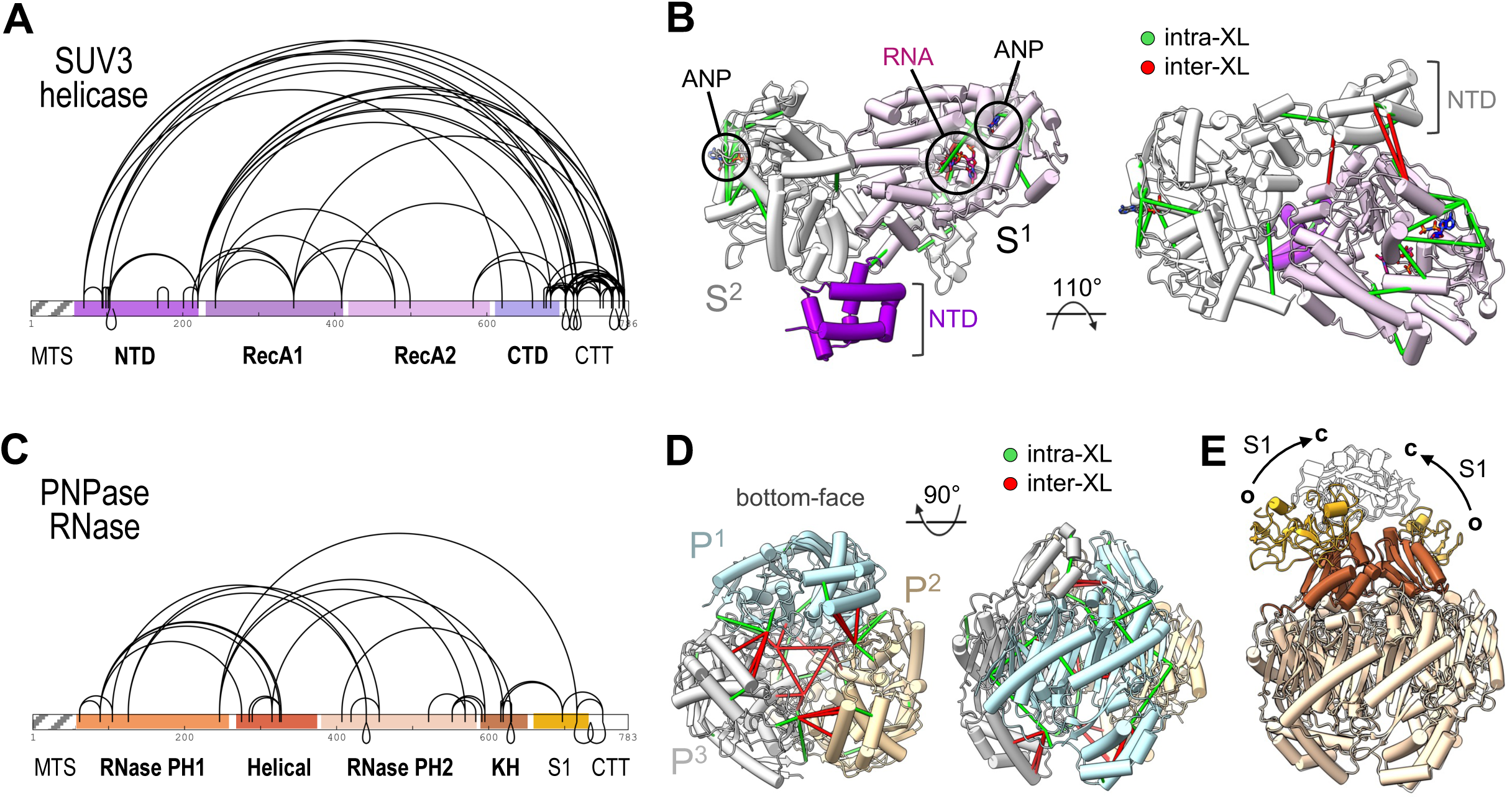
Crosslinking mass spectrometric analysis of SUV3 and PNPase. *A*, BS3 chemical crosslinks identified within the SUV3 helicase. *B*, Orthogonal views of the asymmetric SUV3 dimer bound to AMPPNP (ANP) and RNA with intramolecular (green) and intermolecular (red) crosslinks (XL) shown as solid lines. *C*, BS3 chemical crosslinks identified within the PNPase exoribonuclease. *D*, Orthogonal views of the homotrimetric PNPase crystal structure (PDB 3U1K, (Lin et al. 2012)), lacking the S1 domains, with the three promoters colored blue, yellow, and grey. Intramolecular (green) and intermolecular (red) crosslinks (XL) are shown as solid lines. *E*, Overlay of the open (o) and closed (c) states of the PNPase homotrimer (PDB 9KJR, 9KJT, (Li et al. 2025)), with black arrows indicating the conformational flexibility of the S1 domains. The mitochondrial targeting signal (MTS) removed for these studies is shown in grey diagonal lines for reference.

Inter-molecular crosslinks between SUV3 and PNPase are consistent with an arrangement where the helicase is poised at the exoribonuclease channel entrance. Two overall patterns emerged from unique crosslinks connecting SUV3 to PNPase (Figure 5A). Multiple crosslinks (total 8) connect the SUV3 CTT to the PNPase barrel, likely representing the broad conformational sampling of the labile helicase tail oppose to a stable interface. This interpretation is supported by our affinity pulldown assay demonstrating that the SUV3 CTT is dispensable for RNA Degradosome assembly. An additional 7 unique crosslinks connect the N-terminus of SUV3 to the C-terminus of PNPase (Figure 5A). We interpret these crosslinks as defining a stable SUV3-PNPase interface, as deletion of either the SUV3 NTD or the PNPase S1 domain abolishes complex formation. We suspect this subset of SUV3-PNPase crosslinks likely represent an in-solution ensemble of conformational states that are supported through a stable SUV3-PNPase interface (Jain et al. 2022; Li et al. 2025; Patra et al. 2026).

**Figure 5.**
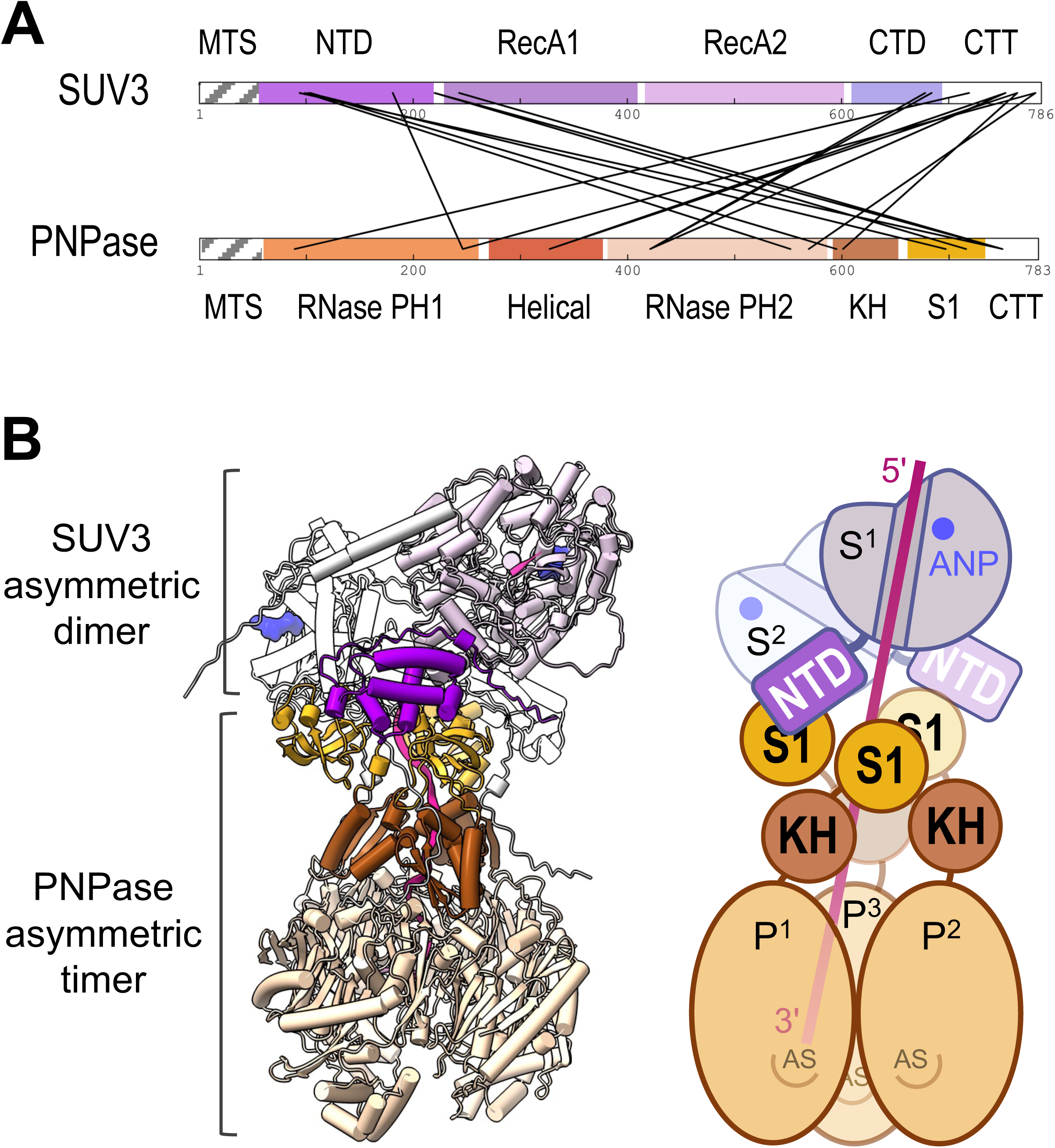
The SUV3 NTD positions the helicase at the pore entrance of the PNPase exoribonuclease. *A*, BS3 chemical crosslinks identified between the SUV3 helicase and the PNPase exoribonuclease. Enriched crosslinks connect the SUV3 NTD domain to the PNPase S1 domain. The mitochondrial targeting signal (MTS) removed for these studies is shown in grey diagonal lines for reference. *B*, A model prediction of the RNA Degradosome complex and corresponding cartoon schematic. AlphaFold predicted the organization of a single SUV3 protomer (purple) bound to the RNA-threaded PNPase homotrimer (orange). The RNA-bound SUV3 protomer from the asymmetric SUV3 dimer (white; PDB 9VD1, (Patra et al. 2026)) was overlaid onto the AlphaFold model to illustrate a possible arrangement of the SUV3-PNPase complex at 2:3 stoichiometry. The overall organization of the model aligns well with the experimental data presented in this work. RNA is shown in pink and AMPPNP nucleotides are shown in blue. AS defines the active site of the exoribonuclease.

### Molecular organization of the RNA Degradosome assembly

To gain structural insight into the SUV3-PNPase interface in the absence of high-resolution experimental data, we generated AlphaFold structure predictions. Model predictions of the hetero-pentameric SUV3-PNPase complex often varied in overall structural organization and were generally low in confidence. Because we demonstrate that SUV3 dimerization is not required for its interaction with PNPase, we generated a structure prediction of a single SUV3 protomer bound to the RNA-threaded PNPase trimer (Supplementary Figure S4). The predicted SUV3 fold is similar to the RNA-bound protomer of the experimentally determined asymmetric SUV3 dimer (r.m.s.d. 0.923 Å) (Supplementary Figure S4). The modeled PNPase barrel also aligns well with both the open and closed cryoEM states, although the relative orientations of the S1 domains differ (Supplementary Figure S4). In the overlay of the asymmetric SUV3 dimer with the modeled RNA Degradosome, the NTD of the RNA-bound SUV3 protomer does not participate in dimerization, leaving it available to interact with the proximal PNPase S1 domain (Figure 5B). Moreover, the polarity of the SUV3 bound RNA is oriented with the 3’-end positioned towards the PNPase channel, allowing for 3’→5’ RNA degradation.

## Discussion

In this study we define the molecular determinants underlying assembly of the human RNA Degradosome. Using complementary chemical crosslinking mass spectrometry and biochemical analyses, we identify the PNPase S1 domain and SUV3 NTD as key determinants of complex formation and show that protein oligomerization is not strictly required. The SUV3-PNPase residue-level constraints refine existing models of the human RNA Degradosome architecture and support a mechanism in which the SUV3 helicase engages the channel entrance of the PNPase exoribonuclease to facilitate processive mitochondrial RNA degradation.

Human and yeast RNA Degradosome complexes employ distinct strategies to couple RNA remodeling with ribonuclease activity. Like humans, yeast mitochondria rely on a helicase-nuclease machinery for bulk RNA turnover, however the yeast SUV3 helicase associates with the structurally distinct 3′→5′ exoribonuclease Dss1 in place of PNPase (Szczesny et al. 2012). The crystal structure of the *Candida glabrata* SUV3-Dss1 complex reveals a minimal architecture in which a single SUV3 protomer associates with the channel entrance of the monomeric Dss1 ribonuclease (Razew et al. 2018). Although Dss1 contains an S1 domain and forms a narrow central channel for unstructured RNA threading and degradation, the S1 domain does not mediate SUV3 association in contrast to human PNPase (Razew et al. 2018). Human and yeast SUV3 share a conserved domain organization comprising an NTD, tandem RecA1 and RecA2 domains, and a CTD, but their NTDs differ in both sequence and structure (Jedrzejczak et al. 2011; Razew et al. 2018; Patra et al. 2026). Moreover, in human SUV3, the NTD is connected to the helicase core by a helix restricting its relative position, whereas the yeast NTD is connected by a flexible linker and is highly mobile (Razew et al. 2018). Consistent with these differences, the yeast SUV3 NTD does not contact the Dss1 ribonuclease nor is it required for RNA degradation (Razew et al. 2018). Together, this suggests that the interface formed by the human SUV3 NTD and PNPase S1 domain is a distinct mode of helicase-nuclease coupling.

The Human RNA Degradosome assembles through an interface that is compatible with, but not dependent upon protein oligomerization. We demonstrate that a single PNPase moiety comprising the KH and S1 accessory domains and C-terminal tail is sufficient for an SUV3 association. Likewise, disrupting SUV3 homodimerization by removing its C-terminal tail has no appreciable effect on PNPase binding. Although the domains mediating the helicase-nuclease interaction in the mitochondrial RNA Degradosome are evolutionarily distinct, the ability of individual protomers to associate is consistent with the 1:1 stoichiometry of the yeast SUV3-Dss1 complex (Razew et al. 2018). Therefore, protein oligomerization may stabilize or regulate the helicase-nuclease interface to enhance RNA degradation without being explicitly required for supporting the SUV3-PNPase interaction.

Positioning SUV3 at the PNPase channel entrance may facilitate coordinated RNA remodeling and degradation. Here, we propose a previously uncharacterized PNPase binding function of the SUV3 NTD, suggesting that the NTD has dual roles within the asymmetric SUV3 dimer. The NTD of the auxiliary protomer contributes to SUV3 homodimerization to support RNA processivity whereas the NTD of the catalytic protomer likely engages the PNPase channel entrance to tightly couple SUV3 helicase activity with PNPase-mediated RNA degradation (Patra et al. 2026). This proposed molecular architecture is supported by data presented in this work and provides context for the previously reported SUV3 CTD deletion that disrupts PNPase binding, as the deleted position is near the channel entrance in the auxiliary SUV3 protomer (Wang et al. 2009). By using domains that are not conserved across the broader SF2 helicase superfamily, the SUV3 NTD and CTD may provide molecular handles for anchoring the helicase to PNPase without constraining the extensive rachet-like conformational changes of its RecA domains during nucleotide binding, hydrolysis, and release that drive RNA remodeling and translocation (Gu and Rice 2010). The conformational flexibility of the S1 domains may further accommodate important dynamic movements across the helicase-nuclease machinery while maintaining a stable SUV3-PNPase interaction. Collectively, this work provides a mechanistic understanding for how the human RNA Degradosome couples SUV3-mediated RNA remodeling and translocation with substrate degradation by the PNPase ribonuclease to maintain mitochondrial RNA homeostasis.

## Materials and Methods

### Molecular Cloning of Bacterial Expression Plasmids

Bacterial expression plasmid (pMP536) of codon-optimized *Homo sapiens* (Hs) SUV3 (residues 46-786) with an upstream 6xHis-TEV sequence was provided by GenScript and inserted into pETDuet-1 between NcoI/BamHI. GenScript subsequently created a bacterial PNPase and SUV3 co-expression plasmid (pMP542) by inserting codon-optimized tagless HsPNPase (residues 47-783) between NdeI/EcoRV of pMP536. Residues 1-46 were omitted from PNPase and 1-45 from SUV3 to mimic the post-proteolytic protein variants generated upon the removal of the mitochondrial targeting signal (MTS). For *in vitro* RNA degradation assays, tagged HsPNPase (residues 47-783) was amplified from pMP542 to include an upstream 6xHis-TEV sequence and inserted into pHis2 (Sheffield et al. 1999) between BamHI/XhoI to generate pMP852. All plasmids were verified by DNA sequencing. Please refer to Table S1 for a list of all bacterial expression plasmids used in this study.

### Molecular Cloning of Mammalian Expression Plasmids

HsPNPase (pMP662) and HsSUV3 (pMP660) cDNAs were provided by Origene in pCMV6-entry vectors between SgfI/MluI. Full-length PNPase (residues 1-783) was PCR amplified from pMP662 and inserted between SgfI/MluI of a pCMV6-entry_3xFlag modified vector were the 3xFlag sequence (DYKDHDGDYKDHDIDYKDDDDK) was added between MluI/PmeI (pMP904). Full-length SUV3 (residues 1-786) was PCR amplified from pMP660 and inserted between SgfI/MluI of a pCMV6-entry_TwinStrep modified vector were the TwinStrep sequence (GSAWSHPQFEKGGGSGGGSGGSAWSHPQFEK) was inserted between MluI/PmeI (pMP899). PNPase and SUV3 variants were generated by Q5 site-directed mutagenesis (NEB) using templates pMP904 and pMP899, respectively. All PNPase and SUV3 variants retained their predicted upstream MTS which are harbored within residues 1-43 and residues 1-77, respectively. A 3-fold symmetric control scaffold (^3x^CTL) was engineered by amplifying HsPCNA (residues 1-261) using Addgene plasmid 58043 as a template and inserted the PCNA amplicon downstream of the human PNPase MTS (residues 1-43) by overlap PCR. To display three copies of the human PNPase C-termini on the 3-fold control scaffold, the human PNPase KH-S1-tail (residues 599-783) sequence was inserted downstream of PCNA (^3x^PCT). A 2-fold symmetric scaffold (^2x^CTL) was created by GenScript by codon optimization of the *Escherichia coli* (Ec) β clamp (residues 1-366) which was inserted downstream of the human PNPase MTS (residues 1-43). The human PNPase C-termini were displayed on the 2-fold control scaffold by inserting the human PNPase KH-S1-tail (residues 599-783) sequence downstream of the β clamp (^2x^PCT). An asymmetric 1-fold symmetric control scaffold (^1x^CTL) was engineered by GenScript by synthesizing a construct consisting of the Ec β clamp (residues 1-366) fused via a glycine-serine linker (GGGGSGGGGSGGGGS) to a second Ec β clamp (residues 1-366), positioned downstream of the human PNPase MTS (residues 1-43). To display the PNPase C-terminus, the human PNPase KH-S1-tail (residues 599-783) sequence was inserted upstream of the stop codon (^1x^PCT). All plasmids were verified by DNA sequencing. Please refer to Table S2 for a list of all mammalian expression plasmids used in this study.

### Protein Expression and Purification

Recombinant human SUV3 was produced as described previously (Van Riper et al. 2025). Recombinant human PNPase was overexpressed in *E. coli* C41(DE3) cells (MilliporeSigma), induced with 0.5 mM isopropyl β-D-thiogalactopyranoside and incubated overnight at 18°C. Cells harboring recombinant PNPase were resuspended in lysis buffer (50 mM Tris pH 8.0, 50 mM K_2_HPO_4_, 300 mM KCl, 25 mM imidazole, 2 mM 2-mercaptoethanol, 1% Triton-X100, 10% glycerol). Cells were lysed by sonication and clarified at 26,916 x g for 50 min at 4°C. Clarified lysate was applied to a gravity flow column loaded with His60 Ni superflow resin (Clontech), washed with lysis buffer supplemented with 2 M KCl, and eluted with lysis buffer supplemented with 300 mM imidazole. PNPase was subsequently resolved over a HiLoad 16/600 Superdex-200 Prep Grade (Cytiva) gel filtration column equilibrated with storage buffer (25 mM potassium phosphate pH 8, 100 mM KCl, 0.5 mM EDTA, 2 mM MgCl_2_, 5% glycerol). PNPase variants were further purified by a HiTrap Heparin HP column equilibrated with heparin buffer (25 mM potassium phosphate pH 8, 5 mM MgCl_2_, 5% glycerol) supplemented with 150 mM NaCl. PNPase was eluted with a linear gradient from 150 mM to 500 mM NaCl and resolved over a Superose 6 Increase 10/300 GL gel filtration column equilibrated with storage buffer. PNPase was concentrated using a 10,000 NMWL centrifugal filter (Millipore Sigma) and flash frozen for long-term storage at –80°C.

### Degradosome Complex Assembly

Purified PNPase (3.75 μM), SUV3 (2.5 μM), or PNPase-SUV3 mixtures (3.75 μM:2.5 μM) were dialyzed in 10K MWCO Slide-A-Lyzer mini dialysis devices (ThermoFisher Scientific) overnight at 4°C in dialysis buffer (25 mM potassium phosphate pH 8, 80 mM NaCl, 2 mM MgCl_2_, 2 mM 2-mercaptoethanol, 5 mM ATP, 15% glycerol). Protein samples were concentrated using Amicon Ultra-0.5 centrifugal filter units (Millipore Sigma).

### Chemical Crosslinking Mass Spectrometry

Crosslinked RNA Degradosome mixture was analyzed by mass spectrometry as described previously (Pillon et al. 2019) with minor modifications. Briefly, the Degradosome (10 μM) was prepared in storage buffer and incubated for 5-minutes in the absence or presence of 3, 6, 12, 24, 48, 96, 192, or 384 μM bis(sulfosuccinimidyl)suberate (BS3; Sigma) and analyzed on a denaturing polyacrylamide gel. A final concentration of 96 μM BS3 produced multiple crosslinked species with minimal sample smearing and was subsequently used for follow-up analysis. The reaction was quenched with 30 mM Tris pH 7.5 for 15 minutes and digested with equal volume of trypsin (Promega) at 0.1 μg/μl in 50 mM ammonium bicarbonate (pH 7.9) overnight at 37 °C and stored at −80 °C for subsequent MS analysis. Protein digests were analyzed using LC/MS with a Q Exactive Plus mass spectrometer (ThermoFisher Scientific) interfaced with an M-Class nanoAcquity UPLC system (Waters Corporation) using a trap and elute strategy including a 75 μm × 150 mm HSS T3 C18 analytical column (1.8 μm particle, Waters Corporation) and a 180 μm × 20 mm Symmetry C18 trap column (5 μm particle, Waters Corporation). Putative crosslinked peptides were identified using Protein Prospector (UCSF). The data were searched against the sequences of the recombinant proteins and included trypsin specificity with up to 3 missed cleavages. Variable modifications included methionine oxidation, deamidation, incorrect monoisotopic assignment, and DSS crosslinking. The putative crosslinked peptide output was triaged by limiting the mass error of putative crosslinks to 3 standard deviations from the average error (3X STD was 6.4 ppm for this data set); requiring a Score Difference value >or = to 3.0, except for the cases where the peptide was duplicated, or the peptide was < or = to 6 amino acids in length; and total expectation values were below 1 × 10^−4^.

### In vitro RNA Cleavage Assay

Ribonuclease activity was measured as described previously (Wang et al. 2009) with minor modifications. Briefly, a 30-nucleotide fluorescein (FI) labeled 5′-FI-GUUGAGAGAGAGA GAGUUUGAGAGAGAGAG-3′ RNA was annealed to a 22-nucleotide 5′-CUCAAACUCUC UCUCUCUCAAC-3′ RNA to form a 3′-overhang substrate. Protein variants (2 μM) were incubated with RNA (50 nM) in dialysis buffer supplemented with 5 mM ATP at 37°C between 1-20 minutes, unless indicated otherwise. Reactions were stopped with equal volume urea loading dye and incubated with 25 mM EDTA and 1 mg/mL proteinase K (NEB) for 15 minutes at room temperature. Samples were resolved by 15% TBE-urea PAGE with 1x Tris-borate-EDTA buffer. Gels were visualized using a Typhoon 5 (Cytiva) or ChemiDoc MP (Bio-Rad). All *in vitro* RNA cleavage assays were performed in triplicate and representative gels are show in the figures.

### Co-Immunoprecipitation and Western Blot Analysis

Plasmid transfections were performed with 20-40 mL cultures of HEK293F cells (Thermo Fischer Scientific). Cells were transfected with 1 μg/mL of purified plasmid DNA and 2 μg/mL of polyethylenimine (Polysciences Inc.). Cells were harvested 48 hours after transfection and stored at –80°C. Cells were lysed in 25 mM potassium phosphate pH 8, 150 mM NaCl, 2 mM MgCl_2_, 1% Triton X-100, 5% glycerol, 0.25 U/μL benzonase, 1 mM benzamidine, 1 mM PMSF, 5 μg/mL leupeptin, and 0.7 μg/mL pepstatin A with gentle rocking for 30 minutes at 4°C. Lysate was clarified at 21,300 x g for 20 minutes at 4°C and incubated with Strep-Tactin XT 4Flow high capacity resin (IBA Life Sciences) or MagStrep “type3” XT beads (IBA Life Sciences) for 1 hour at 4°C. Resin was washed four times with 250 μL of wash buffer (25 mM potassium phosphate pH8, 150 mM NaCl, 2 mM MgCl_2_, 5% glycerol, 0.05% Tween-20), twice with 250 μL of wash buffer supplemented with 10 mM ATP, and one time with 250 μL wash buffer. Samples were resolved by SDS-PAGE and transferred to PVDF membrane using the Trans-Blot Turbo RTA Transfer kit (Bio-Rad) and Trans-Blot Turbo transfer system (BioRad). Membranes were block for 1 hour using 5% nonfat milk in 1x TBST and immediately incubated overnight at 4°C with primary antibody prepared in 5% nonfat milk, 1% (w/v) BSA in 1x TBST. Membranes were incubated with anti-Strep-tag II mouse monoclonal primary antibody (EMD Millipore, 71590-3, 1:1000) for detection of Twin-strep tagged SUV3 variants and anti-flag rabbit primary antibody (Sigma, F7425, 1:1000) for visualization of 3xFlag tagged PNPase variants. Membranes were washed three times in 1x TBST and incubated for 1 hour with goat anti-mouse IgG HRP conjugate secondary antibody (EMD Millipore, AP127P, 1:1000) or goat anti-rabbit IgG HRP conjugate secondary antibody (Jackson ImmunoResearch Lab., 111-035-003, 1:2000). Membranes were wash three times with 1X TBST prior to adding enhanced chemiluminescence detection reagent (Advansta) and imaging the blots on a ChemiDoc MP imaging system (Bio-Rad). All co-immunoprecipitation experiments were performed in triplicate and representative blots are show in the figures.

## Acknowledgements

We thank the members of the Pillon lab for their valuable feedback on the manuscript content and figure design. We are grateful to Dr. Robin Stanley for access to liquid chromatography instrumentation. This work was supported by the US National Institute of Health Extramural Research Program, National Institute of Environmental Health Sciences (NIEHS; R00ES030735 to M.C.P.) and the US National Institutes of Health grant R35GM147123 (MCP) issued to Baylor College of Medicine and transferred to the State University of New York at Buffalo. This research was supported in part by the Intramural Research Program of the NIH, National Institute of Environmental Health Sciences (J.G.W. ES102488). The contributions of the NIH author(s) are considered Works of the United States Government. The findings and conclusions presented in this paper are those of the author(s) and do not necessarily reflect the views of the NIH or the U.S. Department of Health and Human Services.

## Data availability

All data is included in the manuscript figures and supplemental information. The MS data has been deposited in the ProteomeXchange Consortium via the PRIDE partner repository with the dataset identifier PXD083493. All corresponding raw data files will be made available upon request to the corresponding author, Monica C. Pillon.

