## Supplementary Materials for "Molecular architecture and assembly of the human RNA Degradosome"

**This PDF includes:**

Supplemental Figure S1 to S4
Supplemental Table S1 to S2

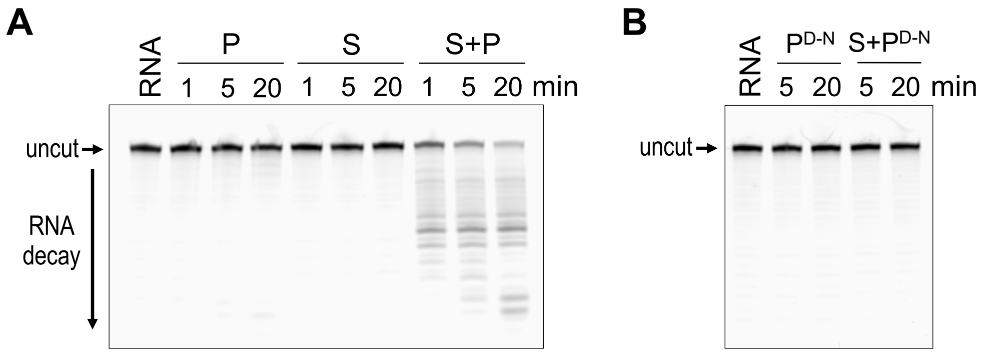


**Figure S1** Reconstitution of substrate degradation by the RNA Degradosome. *A*, RNA degradation activity of human PNPase (P), SUV3 (S), and the RNA Degradosome (S+P) variants (2 μM) with 3′-overhang RNA (50 nM) over a time course. *B*, RNA degradation activity of the human PNPase catalytic variant D544N in the absence ($P^{D\text{-}N}$) and presence of SUV3 ($S+P^{D\text{-}N}$) with 3′-overhang RNA (50 nM) over a time course. Representative gels are shown where sample "RNA" is an RNA alone control. Horizontal arrows mark intact substrate (uncut) and a vertical arrow marked "RNA decay" defines the migration of multiple RNA products.

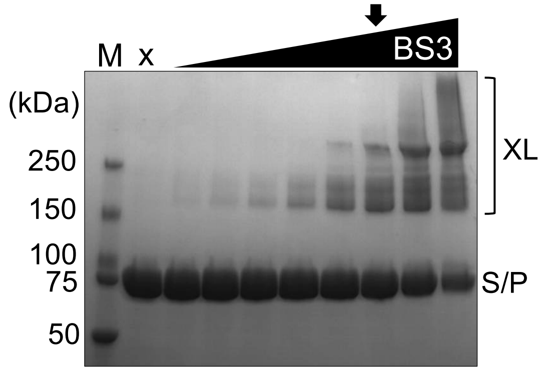


**Figure S2** Chemical crosslinking of the RNA Degradosome sample. The SUV3 (S)-PNPase (P) mixture (10 µM) was incubated with chemical crosslinker bis(sulfosuccinimidyl)suberate (BS3) at a final concentration of 3, 6, 12, 24, 48, 96, 192, and 384 µM for 5 minutes at 22 °C. The reaction was resolved by SDS-PAGE analysis where the migration of crosslinked (XL) species are marked by a bracket. A non-crosslinked control is defined by “X” and a black arrow identifies the condition used for subsequent mass spectrometry analysis.

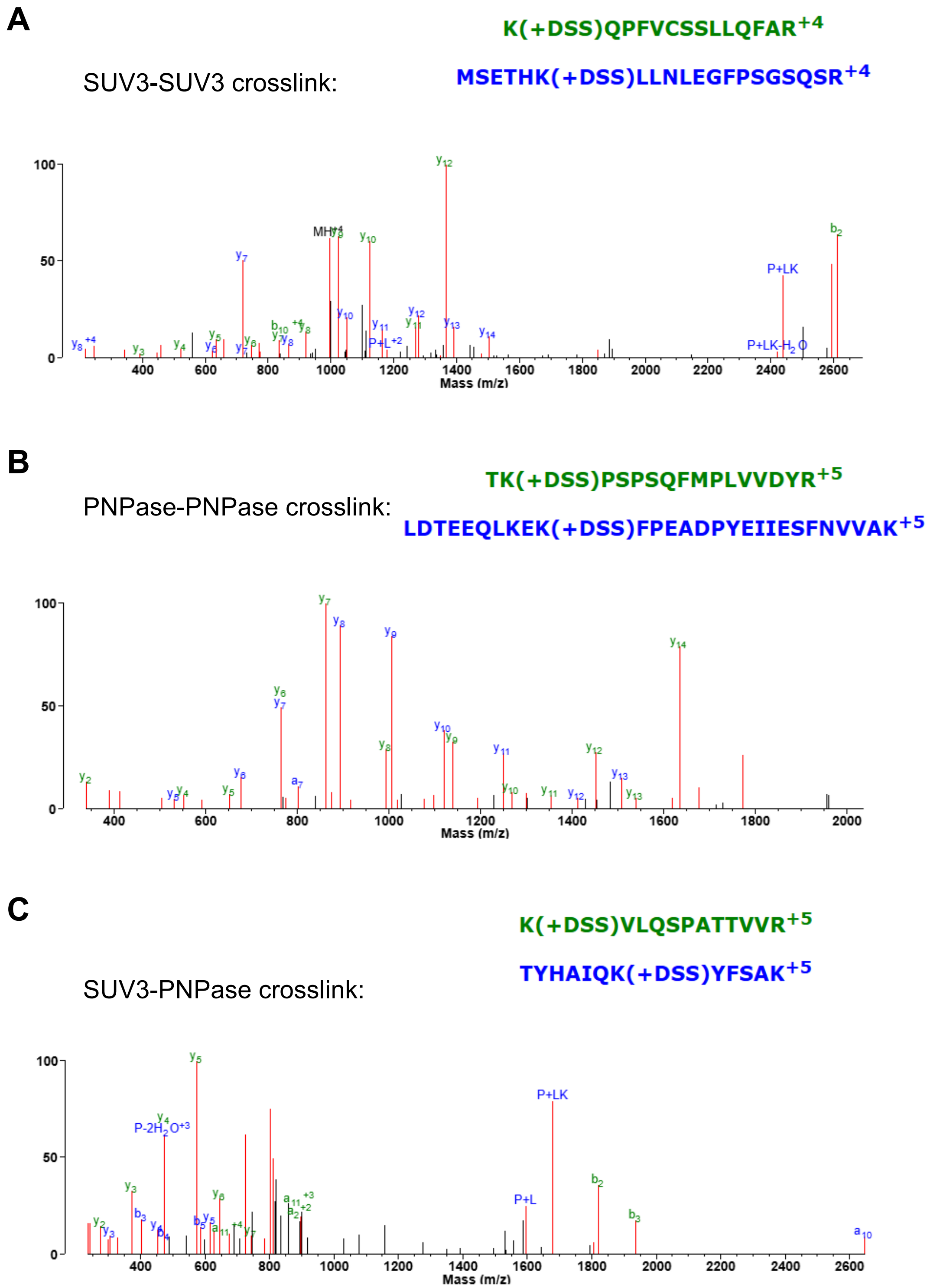


**Figure S3** Representative tandem mass spectrometry spectra. *A*, SUV3-SUV3 crosslink, *B*, PNPase-PNPase crosslink, and *C*, SUV3-PNPase crosslink. Crosslinked peptides are shown at the top of each spectra with "+DSS" indicating the crosslink amino acid.

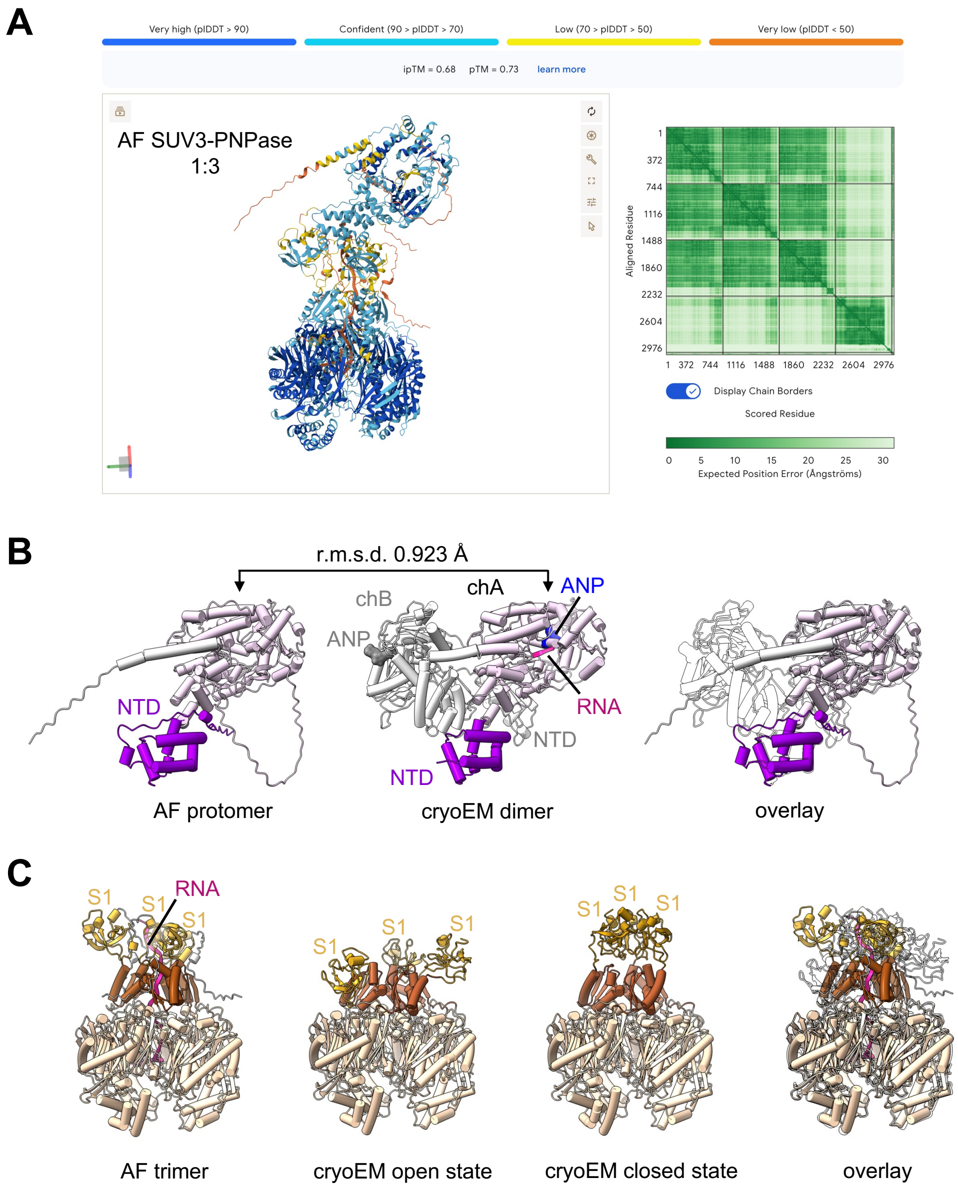


**Figure S4** Model prediction of the RNA Degradosome and its subunits. *A*, Cartoon representation of an AlphaFold (AF) model prediction of an SUV3 protomer bound to the PNPase homotrimer threaded with a 26-nucleotide model RNA substrate (5′-GAGAGAGAGAGAGAGAGAGAG AGAGA-3′). *B*, Cartoon representation of an AF model prediction of full-length SUV3 and the SUV3 asymmetric homodimer bound to AMPPNP and single-stranded RNA (PDB 9VD1, (Patra et al. 2026)). An overlay of the models is shown on the right where the experimental structure is colored white. *C*, Cartoon representation of an AF model prediction of full-length homotrimeric PNPase, the PNPase open state (PDB 9KJR, (Li et al. 2025), and the PNPase closed state (PDB 9KJT, (Li et al. 2025)). An overlay of all the models is shown on the right.

**Table S1.** *E. coli* expression plasmids of Human PNPase and SUV3 used in this study

| Plasmid | Description | Vector | Source |
|---|---|---|---|
| pMP542 | PNPase+SUV3;<br>tagless HsPNPase, residues 47-783,<br>6xHis-TEV-HsSUV3, residues 46-786 | pET-Duet1 | GenScript |
| pMP584 | PNPase WT;<br>HsPNPase-6xHis, residues 47-783 | pST39 | This study |
| pMP852 | PNPase WT;<br>6xHis-TEV-HsPNPase, residues 47-783 | pHis2 | This study |
| pMP884 | PNPase catalytic mutant (D-N);<br>6xHis-TEV-HsPNPase, residues 47-783 D544N | pHis2 | This study |
| pMP536 | SUV3 WT;<br>6xHis-TEV-HsSUV3, residues 46-786 | pET-Duet1 | (Van Riper et al. 2025) |

**Table S2.** Mammalian plasmids used in this study

| Plasmid | Description | Vector | Source |
|---|---|---|---|
| pMP662 | PNPase 1-783; HsPNPase-Myc_DDK, residues 1-783 | pCMV6-entry | Origene |
| pMP904 | PNPase 1-783; HsPNPase-3xFlag, residues 1-783 | pCMV6-entry_3xFlag | This study |
| pMP909 | PNPase 1-753; HsPNPase-3xFlag, residues 1-753 | pCMV6-entry_3xFlag | This study |
| pMP910 | PNPase 1-673; HsPNPase-3xFlag, residues 1-673 | pCMV6-entry_3xFlag | This study |
| pMP911 | PNPase 1-598; HsPNPase-3xFlag, residues 1-598 | pCMV6-entry_3xFlag | This study |
| pMP980 | $^{1x}$CTL; HsPNPase, residues 1-43, fused to EcSliding clamp-linker- EcSliding clamp'-3xFlag, residues 1-366, linker, 1-366 | pCMV6-entry_3xFlag | This study |
| pMP970 | $^{1x}$PCT; HsPNPase, residues 1-43, fused to EcSliding clamp-linker- EcSliding clamp', residues 1-366, linker, 1-366, fused to HsPNPase-3xFlag, residues 599-783 | pCMV6-entry_3xFlag | This study |
| pMP951 | $^{2x}$CTL; HsPNPase, residues 1-43, fused to EcSliding clamp-3xFlag, residues 1-366 | pCMV6-entry_3xFlag | This study |
| pMP950 | $^{2x}$PCT; HsPNPase, residues 1-43, fused to EcSliding clamp, residues 1-366, fused to HsPNPase-3xFlag, residues 599-783 | pCMV6-entry_3xFlag | This study |
| pMP936 | $^{3x}$CTL; HsPNPase, residues 1-43, fused to HsPCNA-3xFlag, 1-261 aa | pCMV6-entry_3xFlag | This study |
| pMP929 | $^{3x}$PCT; HsPNPase, residues 1-43, fused to HsPCNA, residues 1-261, fused to HsPNPase-3xFlag, residues 599-783 | pCMV6-entry_3xFlag | This study |
| pMP660 | SUV3 1-786; HsSUV3-Myc_DDK, residues 1-786 | pCMV6-entry | Origene |
| pMP899 | SUV3 1-786; HsSUV3-TwinStrep, residues 1-786 | pCMV6-entry_TwinStrep | This study |
| pMP919 | SUV3 ΔCTT; HsSUV3-TwinStrep, residues 1-685 | pCMV6-entry_TwinStrep | This study |
| pMP918 | SUV3 ΔNTD; HsSUV3-TwinStrep, residues 1-77, 170-786 | pCMV6-entry_TwinStrep | This study |
